# Machine characterization of rat toxin responses identifies disease states, tolerance mechanisms and organ to whole-body communication

**DOI:** 10.1101/322446

**Authors:** Kenichi Shimada, Timothy J. Mitchison

## Abstract

**Background:** Living organisms are constantly exposed to toxic xenobiotics and have therefore evolved protective responses. In mammals, the liver and kidney play central roles in protecting the organism from xenobiotics, and are at high risk of xenobiotic-induced injury. Liver and kidney damage by drugs and industrial toxins have been extensively studied from both classical histopathologic and biochemical perspectives.

**Methods and Findings:** We introduce a machine learning approach for the analysis of toxicological response. Unsupervised characterization of physiological and histological changes in a large toxicogenomic dataset revealed nine discrete toxin-induced disease states. Transcriptome analysis showed that some of the machine-identified disease states correspond to known pathology, and to known effects of certain toxin classes, but others were novel. Analysis of dynamics revealed transitions between disease states at constant toxin exposure, mostly in the direction of decreased pathology, which implies induction of tolerance. Tolerance correlated with induction of known xenobiotic defense genes and novel decreased ferroptosis sensitivity biomarkers. These data reinforce emerging evidence that ferroptosis drives organ pathology, and suggest that its downreagulation may promote tolerance and recovery. Lastly, mechanism of body weight decrease, a known primary marker for toxicity, was investigated. Combined analysis of food consumption, body weight, and molecular biomarkers indicated that organ disease states promote cachexia by whole-body signaling through Gdf15 and Igf1, suggesting strategies for therapeutic intervention that may be broadly relevant to human disease.

**Conclusions:** Application of machine learning to systematic data collection of physiology, histopathology, transcriptome reveals multiple disease states, tolerance mechanisms and organ to whole-body communication.

## Introduction

A major function of the liver and kidneys is to take up, metabolize and excrete xenobiotics that gain access to the blood. In executing these functions, these organs are at high risk for toxin-induced damage. Liver and kidney toxicities are major concerns in the safety of pharmaceuticals, industrial chemicals and environmental toxins *[1,2]*. Predictive toxicology aims to identify molecular events that precede and cause tissue injury to inform exposure limits and development of less-toxic alternatives *[3,4]*. Toxicogenomics, i.e. collection of transcriptomic and other systematic data across sets of reference toxins in model organisms, is a relatively recent innovation with high potential for improving the mechanistic understanding of toxicities *[5,6]*. Here, we use the Open TG-GATEs database, which collates high-quality transcriptome, histopathology, blood chemistry, tissue and body weight data following administration of 160 chemicals in rats *[7,8]*. These data have previously been analyzed using genecentric classification schemes with the goal of improving predictive toxicology [9–11]. Our approach is instead disease-centric, with more clinically oriented goals: to learn the number and nature of discrete disease states induced by toxins, how the liver and kidney respond to oppose induction of local pathology, and how they also orchestrate organism-wide responses to toxin exposure. We view this approach as conceptually similar to that of physicians seeking to classify the mechanisms of diseases more generally, and we ask, to what extent does unsupervised machine learning discover the same disease-states as physicians?

Supervised classification, which dominates toxicogenomics, seeks to best separate multiple experimental outcomes (especially mRNA expression patterns) into pre-determined phenotypes (e.g., fibrosis or carcinogenicity in the liver) *[12,13]*. It does not test whether the data best support those pre-determined phenotypes versus others, or how multiple phenotypes relate to each other and interconvert *[14,15]*. We therefore sought first to identify mutually exclusive disease states in a data-driven manner, and only then to decipher state-specific molecular mechanisms or biomarkers. To do this, we chose to begin with information a clinician can access in man, rather than classifying transcriptomes. Although transcriptomes offer large amounts of data and the potential for molecular pathway identification *[10,11]*, they are not part of standard clinical diagnosis, and their relationship to disease states is unclear. As a machine-mimic of clinical diagnosis, we started by clustering conditions that exhibit abnormal physiology (*i.e*., blood chemistry and body and tissue weights) and histopathology from the Open TG-GATEs dataset. These data collected in rats model a standard set of clinical measurements applied by physicians to patients with almost any unidentified disease. Using unsupervised clustering, we identified nine discrete disease states that were independently supported by physiological and histopathological data. We then performed a supervised analysis of gene expression data through the lens of these machine-identified disease states. Our combined data analysis revealed that some machine-identified disease states correspond to known disease states and known mechanisms of toxin action but others seem novel. We identified temporal transitions between disease states that provide evidence for the induction of tolerance, and we found distinct gene expression signatures that correlated with tolerance. These included changes in the expression of xenobiotic metabolism genes, as expected, and also novel biomarkers for protection from ferroptosis, a specific form of cell death mediated by runaway lipid oxidation *[16]*. Finally, we explored the role of the liver in mediating whole-organism responses to xenobiotics. We find evidence that the liver communicates with the rest of the body through specific signaling proteins that likely mediate feeding behavior and weight loss.

## Methods

### Normalization of Open TG-GATEs dataset

#### Open TG-GATEs data acquisition

All but food consumption data were downloaded from the Open TG-GATEs website (http://toxico.nibiohn.go.jp/open-tggates) using RCurl package and parsed with XML package. Food consumption data were downloaded from Life Science Database Archive (https://dbarchive.biosciencedbc.jp/en/open-tggates/data-11.html). An administration of one compound, one dose, and one time point is referred to as “treatment condition” or “condition” throughout the text. There were 3,564 conditions tested in total (160 chemicals, 3 doses, 8 time points). Each condition was tested in biological quintuplicates to collect physiology (body and organ weights, blood cell counts, blood chemistry) and histology (diagnosis based on H&E staining of liver and kidney made by toxicologic pathologists) data; three of the five samples were further tested for liver and kidney microarray data; 3,528 and 975 conditions were tested for liver and kidney transcriptome.

#### Drug treatment information

In the Open TG-GATEs dataset, rats were tested with one of 160 compounds (99 drugs, 55 industry toxins, 6 endogenous signaling molecules or metabolites), at three doses determined for each chemical (Fig. S1, Table S1). After single dose treatment, animals were sacrificed at 3, 6, 9, or 24 hrs; after daily repeated dose treatment, animals were sacrificed on 4, 8, 15, or 29 days *[7]*. Some compounds were tested in single or repeated dose treatments only, but 140 compounds were tested for all eight time points. 97 of which were at same doses, but 43 of which were tested at higher doses in single dose than at repeated dose. 365 conditions (compounds at fixed doses) were scheduled for all eight time point testing, but 14 killed animals at later time points so 351 conditions were tested at eight time points. At the time of sacrifice, physiology (hematology, body and tissue weights) and histopathology of liver and kidney were collected for each of the five animals. Further, 3 of 5 animals representing each condition were collected and subject to liver and kidney microarray.

#### Physiology data normalization

Physiology parameters (blood cell counts, blood biochemistry, and body and tissue weights) were measured for each first subject to normalization for each of 3,564 treatment conditions (compounds, doses, time points). We first averaged five biological replicates of each parameter for each condition. Because each parameter was measured in different units and they were not directly comparable, we normalized the value so that they were comparable to each other. As for normalization, we computed the mean and the interquartile range of 3,564 values of each parameter, subtracted the mean from each value, and divided by the interquartile range.

#### Histopathology curation

Histopathology of liver and kidney from every treated animal was diagnosed by toxicologic pathologists. The information consists of the names of phenotype (necrosis, hypertrophy, etc), topography (periportal, centrilobular, etc), and grade (minimal, slight, moderate, severe). Because some histopathology phenotypes (such as necrosis) were often observed even under vehicle treatments, so they were considered independently of compound treatment. Therefore, we trimmed the histopathology observations so that we work only with observations likely induced by compound treatments. To do this, we stratified the observations into topography and grade. For each topography and grade, we counted the number of rats exhibits the phenotype both induced by vehicles and by compounds, and kept the observations only when the ratio of the counts was more than the ratio of rats used in the project (0.336; there were 5950 and 17685 rats used for control and experiments, respectively), and discarded observations otherwise because they were not likely to be induced by compounds. When a compound treatment induced a phenotype in one of the quintuplicates, we claimed the phenotype was induced by the condition.

#### Transcriptome normalization

Microarray experiments were performed in three biological replicates. All the CEL files from rat liver and kidney data were downloaded from the Open TG-GATEs website. There were 14,143 and 3,905 CEL files for the liver and kidney. The CEL files of the same tissue were handled simultaneously for computing an expression matrix using affy, affyio, BufferedMatrix, BufferedMatrixMethods, rat2302.db packages. Particularly, normalization was performed by robust multiarray analysis of BufferedMatrix.justRMA() function of BufferedMatrixMethods, which log2 transformed the resulting expression profiles. Three biological replicates were averaged to produce an expression profile for each condition, and a profile of the corresponding vehicle treatment was subtracted. This gives expression profiles of 3,528 and 975 conditions in liver and kidney, respectively.

#### Food consumption normalization

For 337 conditions (132 unique compounds), food consumption was measured at nine time points (1, 4, 8, 11, 15, 18, 22, 25, 29 days). For these conditions, food consumption of rats administered with compounds were subtracted from that of rats administered with vehicles.

### Identification of disease states / compute physiology and histology over-representation

#### Computing physiology t-SNE

Using 1-Pearson correlation as distance measure between any pairs of treatment conditions in the physiology space, we first computed a distance matrix across 3,564 conditions. We next set a seed *i* for random number generator (RNG) (*i*=1-100), and ran t-SNE based on the calculated distance matrix using Rtsne() function in Rtsne package, to generate a 2-dimensional coordinate of each conditions on the t-SNE map.

#### Filtering disease-associated conditions

Severity scores were computed by counting co-occurring histology phenotypes for liver and kidney and mapped onto t-SNE map. 2-dimensional density landscape of severity scores were computed using bkde2D() function in KernSmooth package. Severity score is recomputed by estimating the severity score from the 2-dimensional density map using interp.surface() function in fields package. Conditions containing higher severity scores than an arbitrary threshold were considered to be associated with diseases and further selected for disease identification.

#### Clustering for identifying disease states

Conditions with higher severity scores were clustered based on their t-SNE coordinates using density based clustering of applications with noise (DBSCAN). This is achieved by dbscan() function in dbscan package. 100 runs from t-SNE to clustering with different RNG seeds, were summarized by ensemble clustering using cl_consensus() function in clue package. This identified 15 clusters that contain 5 to 203 conditions. To gain robust disease states that are induced by multiple compounds, we discarded smaller clusters composed of fewer than 20 conditions, because we expected that such small clusters do not have strong statistical power due to the small sample size in further transcriptome analysis. We recomputed the memberships and likelihoods to limit our interest to larger clusters with ≥ 20 conditions, and found 9 consensus clusters in total ranging from 37 to 203 conditions (10 to 55 unique compounds). At the same time, 2,723/3,564 conditions were identified a non-disease states.

### Characterization of physiology and histology of 9 DSs

#### Relative severity between liver and kidney

Liver and kidney severity scores for each disease was compared to assess which tissue was more affected in terms of histopathology. Relatively affected tissue was assessed by scatterplot (Fig. 2A, top) as well as log ratio: log10(severity_liver_) – log10(severity_kidney_) (Fig. 2A, bottom).

**Fig 1.**
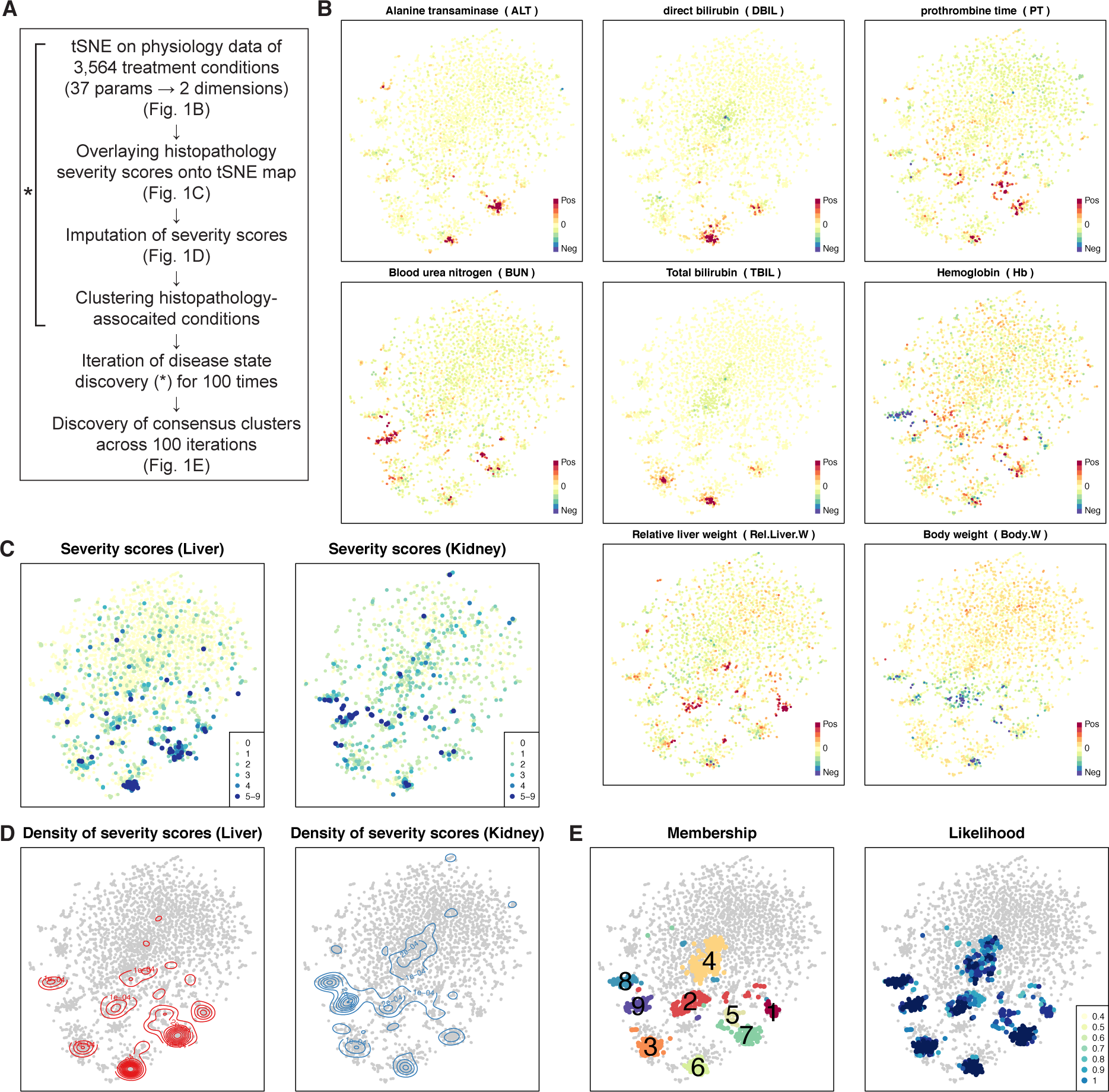
Discovery of 9 disease states using physiology and histopathology data. **(A)** Computational process to discover disease states in Open TG-GATEs. **(B)** Physiology t-SNE map generated by correlation distance and t-SNE. Each point represents the physiology data from one treatment condition. 3,564 conditions were color-coded by intensities of 8 physiology parameters. **(C)** Severity scores of liver and kidney overlaid onto the physiology tSNE map. **(D)** Contour maps indicating densities of severity scores in liver and kidney. **(E)** Membership and likelihood of consistent clusters across DBSCAN on 100 times iterations of tSNE. Numbers on the clusters corresponds to their names, DS1-9, that are consistent throughout the paper. See also Figures S1-6.

**Fig 2.**
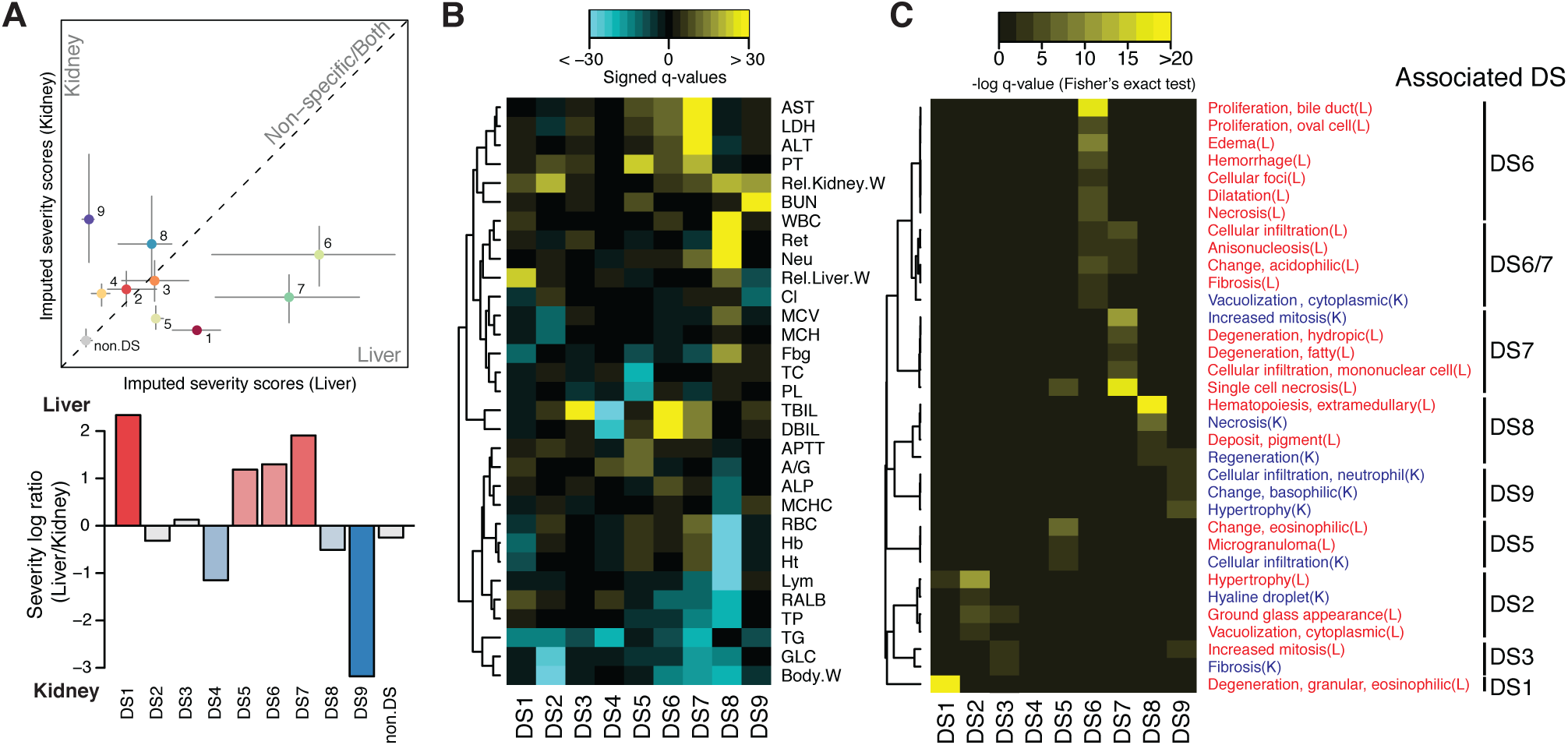
Disease state characterization using blood physiology and histopathology. **(A)** Comparison of liver and kidney severity scores indicate more affected tissue in each DS (top). Severity log ratio of liver over kidney severity scores (bottom). **(B)** Changes in physiology parameters and DSs. Yellow/cyan indicates the parameters are higher/lower than corresponding vehicle treatments in the DS. 31 parameters whose FDR adjusted p-values < 1e-10 (Wilcoxon two-sample test) at least in one DS were shown. **(C)** Histopathology phenotypes and DSs. Relative enrichment of a phenotype among DSs were shown. Yellow indicates more observations in a DS than the others. 34 histopathology phenotypes whose FDR adjusted p-values < 5e-3 (Fisher’s exact test) at least in one DS were shown.

#### Deviation of physiological parameters in each DS

Changes in physiology parameters were assessed by unpaired two-sample two-sided Wilcoxon test between conditions in each DS and conditions in non-DS. Resulting p-values were adjusted to false discovery rate (FDR) (also known as q-values), and further converted to ‘signed log q-values’ *[33]* (Fig. 2B). Physiological parameters whose q-value < 10^−10^ against at least one DS were shown in Fig. 2B.

#### Relative enrichment of histopathological phenotypes among DSs

Among conditions associated with at least one histopathological observation, we assessed if each histopathology phenotype was more observed in a specific DS, using one-sided Fisher’s exact test. All the p-values were FDR-adjusted and converted to singed log q-values, and histopathology phenotypes whose q-values < 5 x 10^−3^ against at least one DS were shown in Fig. 2C.

### Elastic net classification of DS using microarray data

To assess whether liver or kidney transcriptome is powerful enough to distinguish each DS from the rest, we built elastic net classifiers using cv.glmnet() function of glmnet package. For treatment conditions assigned to each DS and the rest of conditions were classified using transcriptome data of either tissue (in total, 3,528 liver or 975 kidney conditions). Both conditions assigned into the DS and not were split into 10 groups randomly (10-fold cross validation), and the elastic net classifier was trained with one of 10 groups being left out, where the conditions were weighted reciprocally proportional to the size of the groups (DS or not). In glmnet(), Binomial family for the response type and area under curve for the type measure were chosen. The left out conditions were predicted using the trained classifier. Cross validation was repeated 10 times, with different ways to divide conditions into 10 groups, and the predicted probabilities were averaged (prevalidation). The performance of the classification was assessed by area under receiver operator curve (AUROC) computed using auc() function of pROC package. To evaluate the performance of classifiers, the same analysis was run on randomly withdrawn conditions with the same sample sizes; AUROC values from randomly withdrawn conditions were substantially lower than classifications of any DSs.

### Pathway analysis

#### Compiling pathways

We assembled 973 pathway information using KEGG.db (v3.2.3), and GO.db (v3.5.0) Bioconductor packages. Using rat2302.db (v3.2.3), org.Rn.eg.db (v3.5.0) packages, we found 914 of which have ≥ 10 genes that were measured in Affymetrix Rat Genome 230 2.0 Array.

#### Computing activity scores

We assess whether the 914 GO and KEGG pathways were activated or inactivated commonly among conditions assigned to each DS, compared to pooled non-DS conditions. To appropriately assess the significance of changes of pathway activity across hundreds or thousands of conditions spanning multiple DSs, we modified gene set enrichment analysis (GSEA) *[49]*. Original GSEA sorts the gene list based on their expressions and one-sample Kolmogorov-Smirnov (KS) test, which assess permutation-based significance of KS statistics. In our method, we first computed KS statistic of each pathway in each condition, and asked if KS statistics of conditions assigned to one DS is overall higher or lower compared to non-DS conditions, using two-sample Wilcoxon test (also known as Mann Whitney *U* test). Resulting p-values were converted to signed log p-values, which we termed “activity scores”. A large positive or negative activity score indicates that a pathway is significantly up- or down-regulated across conditions assigned in the DS compared to non-DS conditions. Note that we decided to not adjust p-values for multiple hypothesis testing for transcriptome analysis because pathways information from two different databases are highly redundant, but instead we chose fairly strict criteria (p = 10^−5^) for calling a pathway’s change significant.

### Transcriptome characterization of DSs

#### DSs similarity based on transcriptome

Using 723 liver and 192 kidney pathways whose transcriptional activity was significantly up or downregulated at least in one DS, we measured similarity of the transcriptome of the 9 DSs using hierarchical clustering, using 1-Spearman correlation as distance measure and a complete linkage method for the clustering (Fig. 3B).

**Fig 3.**
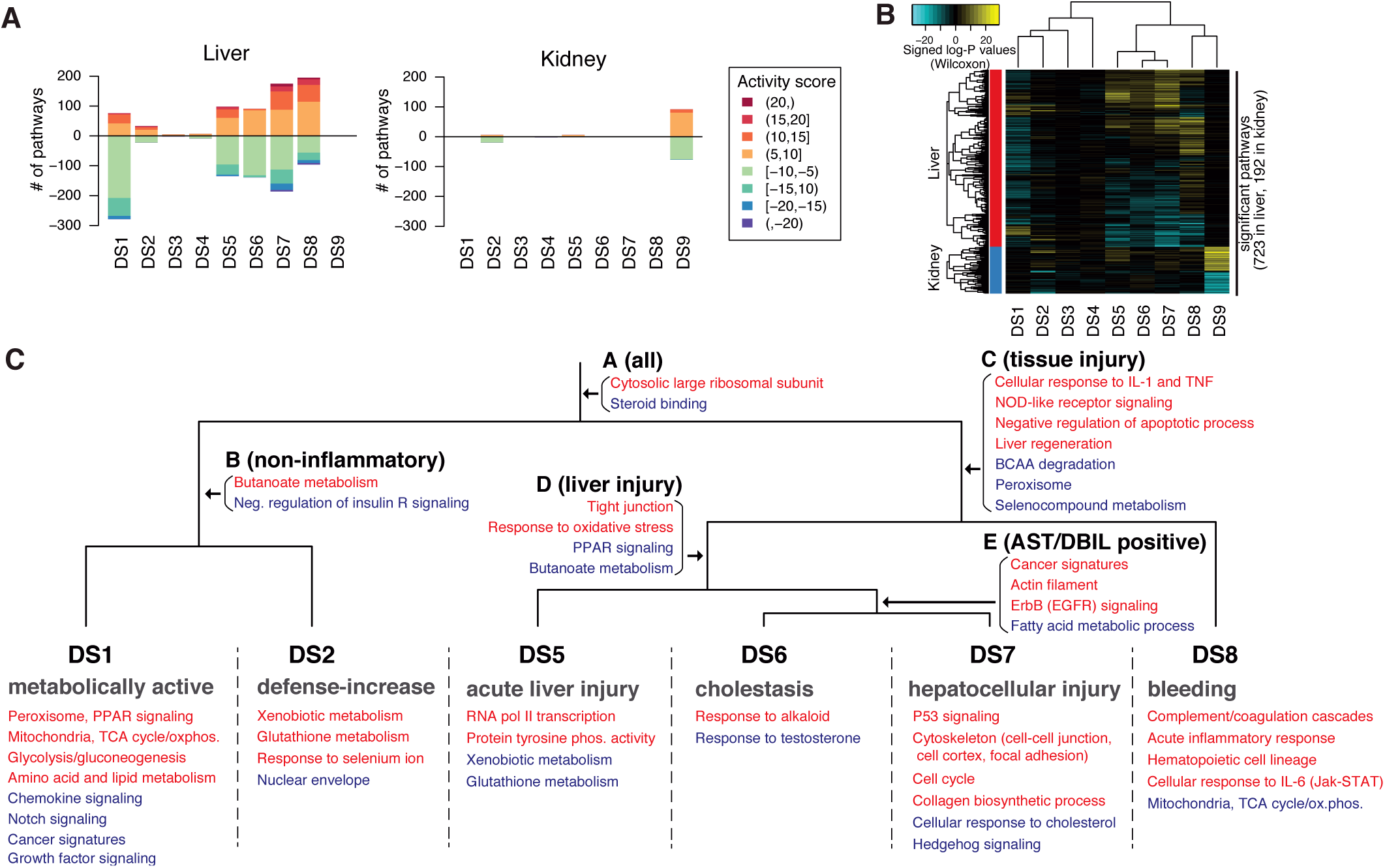
Disease state characterization using transcriptome data. **(A)** Number of significantly up-/down-regulated pathways in each disease state from the liver and kidney transcriptome. Colors of the bars represent pathways with different significance. **(B)** Transcriptional activities of significantly up-/down-regulated pathways in the liver and kidney across all 9 DSs. **(C)** Manually curated map of transcriptional activities of pathways in the liver among six DSs (DS1,2,5-8). Up- and down-regulated are colored in red and blue, respectively. See also Figure S7.

#### DSs characterization among multiple DSs

We mapped 723 pathways in the liver transcriptome based on their activities across six DSs (DS1-2,5-8) whose liver transcriptome was systematically changed. We first checked if each pathway’s change in the liver of the six DSs, comparing their activity score with thresholds (≥ 5 for upregulation, ≤ −5 for downregulation). Then, patterns of up/down-regulations of a pathway were matched with the dendrogram (Fig. S7C-E). Pathways exclusively changed in one direction only in one DS were mapped onto each DS (e.g., xenobiotic metabolism in DS2), pathways commonly changed in multiple DSs were associated to the corresponding branching point in the dendrogram (e.g., cancer signature in DS6-7). In the extreme, a pathway upregulated in all the six DSs (“Large ribosomal subunit” (GO:0042273)) was associated with the top branching point in the dendrogram. There were some pathways that were not mapped to dendrogram. For example, “Terpenoid biosynthesis (rno00900)” was upregulated in three DSs (DS1,2,8), where there were no corresponding points in the dendrogram.

### Disease transition network between DSs

Of the 365 conditions (compounds and doses) scheduled at all eight time points between 3 hr to 29 day (14 of which were scheduled at eight time points but rats were killed before 15-day or 29-day time points, so 351 of 365 were actually tested at eight time points), we first looked at the DS assigned at each time point. 119 conditions did not exhibit any DSs. Of the 246 conditions that took some DSs at least once, 90 took more than one DS across eight time points. In some cases, non-DS states were observed while transitioning from one DS to another, which we removed to highlight the relationship between DSs. The dynamics between DS was visualized using igraph package (Fig. 4B). Transition to and from non-DS (represented by outer open circle) were manually added in Adobe Illustrator.

**Fig 4.**
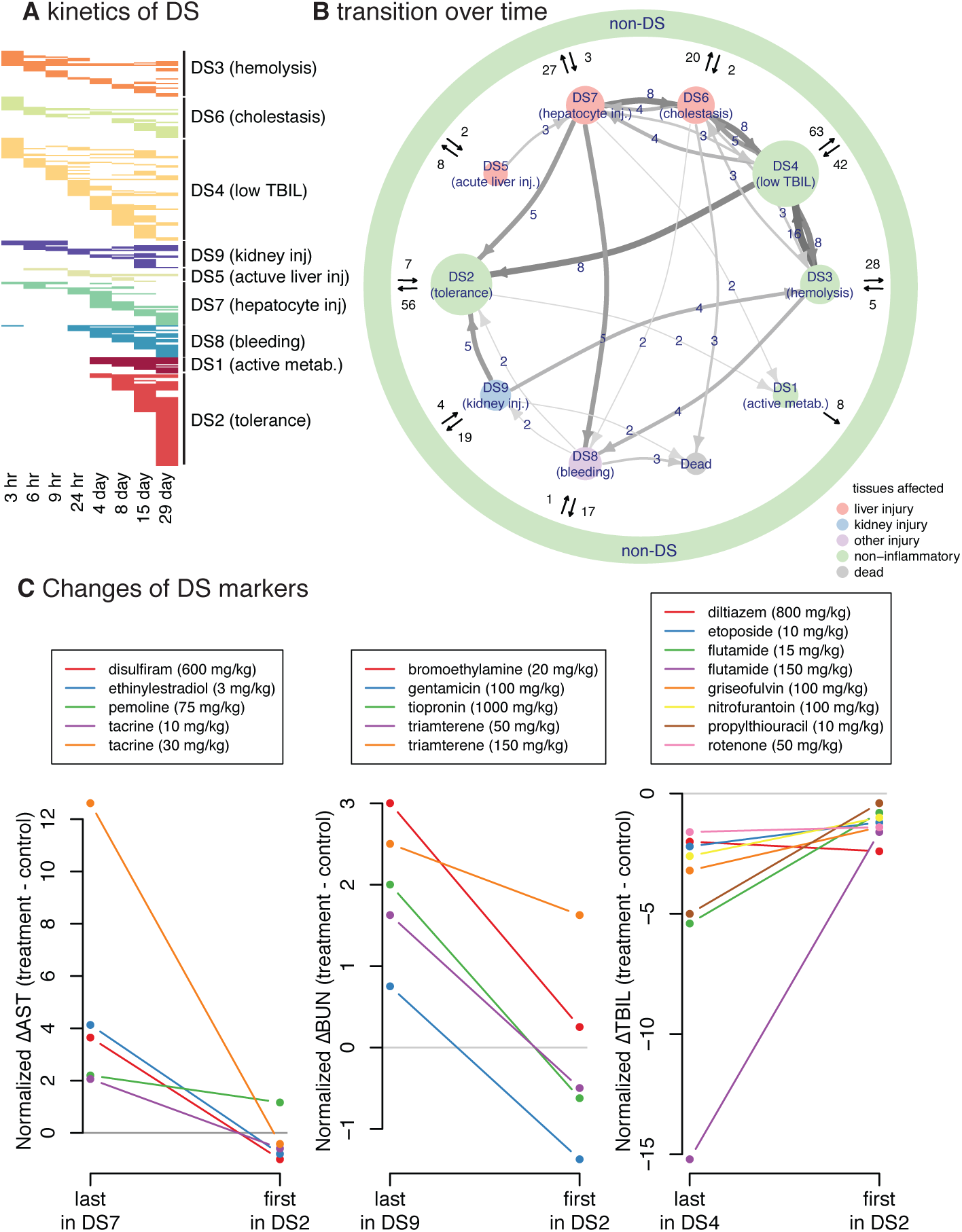
Disease state dynamics. **(A)** Kinetics of DSs, summarizing the frequency and timing of each DS. Each row indicating one treatment (compound and dose) across eight time points. **(B)** Transition between DSs. Size of nodes reflect the number of unique treatments assinged to each DS. Size and color of arrows reflect the number of treatments transitioning between the two nodes. Transitions taken by only one compound were omitted. **(C)** Evidence of tolerance. As for transitions from three DSs to DS2, changes in representative markers for each DS were shown. See also Figure S8.

### Enzyme stratification by co-factors

The 14 pathways upregulated exclusively in DS2 were mostly xenobiotic metabolism enzymes. We evaluated expression of each of these genes in DS2 using one-sample two-sided Wilcoxon test, with resulting p-values converted to singed log p-values. Then, we found the enzymes’ Enzyme Commission (EC) numbers, which was available in org.Rn.eg.db package. All of the enzyme-encoding genes in the 14 pathways were either oxidoreductases (EC1), transferases (EC2), or hydrolases (EC3). They require cofactors for the enzymatic functions. Except for EC3, which requires water as cofactor that is abundant in cells, we regrouped EC1 and EC2 enzymes based on the cofactors: NAD(P)H (EC1.-.1.-, EC1.-.13.-), cytochrome (EC1.-.2.-), oxygen (EC1.-.3.-), disulfide (EC1.-.4.-), flavin (EC1.-.14.-), iron-sulfur (EC1.-.15.-), S-adenosyl methionine (EC2.1.-.-), acyl-CoA (EC2.3.-.-), nucleotide sugar (EC2.4.-.-), glutathione (EC2.5.1.18), ATP (EC2.7.-.-). Finally, the signed log p-values of the enzymes gene expressions were grouped by the co-factors of choice and visualized as boxplot (Fig. 5A). GSEA was performed to see enrichment of NAD(P)H-dependent, GSH-dependent, and all enzyme-encoding gene expressions in DS2 transcriptome.

**Fig 5.**
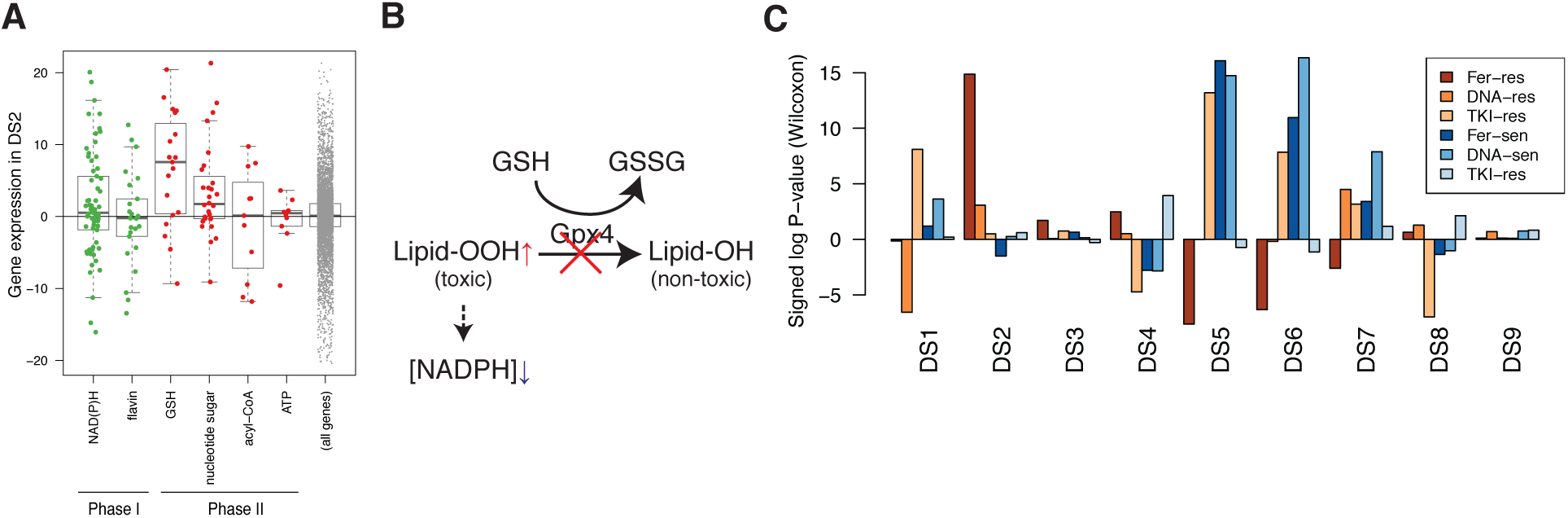
Induced drug tolerance is partly due to resistance to ferroptosis. **(A)** Expression of xenobiotic metabolism enzyme genes grouped by phases of biotransformation and cofactors. Only cofactors with ≥ 10 enzymes were shown. **(B)** Central mechanism of ferroptosis. **(C)** Activity of disease mechanism-specific sensitive and resistant gene signatures in the nine DSs, focusing on three cell death mechanisms (Fer: ferroptosis, DNA: DNA-damaging, TKI:tyrosin kinase inhibition). See also Figure S9.

### Discovery of biomarkers to different cell death phenotypes

We previously showed that 2,565 compounds which were relatively cell-line selectively lethal compounds (i.e., compounds whose GI_50_ values vary across cell lines) in the NCI-60 dataset were classified into 18 mechanistically distinct classes based on the relative growth inhibitory (GI_50_) profiles *[33]*. While most of their mechanisms of action were not yet characterized, a few annotated ones were DNA-targeting compounds (DNA), ferroptosis (Fer), and tyrosine kinase inhibitors (TKI). In the paper, we correlated basal microarray expression profiles with drug sensitivity profiles of each mechanism class, where positive and negative correlations can be interpreted as overexpressed in more resistant or sensitive cell lines, respectively. We extended this approach in this study. We took top 200 most positively and negatively correlated genes in the three cell death phenotypes (DNA, Fer, TKI) as resistant (res) and sensitive (sen) biomarkers for the cell death, because these are genes likely overexpressed in cells resistant or sensitive to each cell death phenotype. Thus, we created 6 human gene sets: DNA-res, DNA-sen, Fer-res, Fer-sen, TKI-res, and TKI-sen. Ensembl’s biomart and Mouse Genome Informatics (http://www.informatics.jax.org/downloads/reports/HOM_AllOrganism.rpt) were used to find 158-185 rat orthologs for each gene set. We computed activity scores of the six gene sets across nine disease states to assess whether they are resistant or sensitive to the three different mechanisms of cell death (Fig. 5C).

### Correlation of pathways with ferroptosis sensitivity signatures

We computed KS statistics of the two curated gene sets, Fer-sen and Fer-res, as well as 914 GO and KEGG pathways. Then computed Spearman correlation between the KS statistics of Fer-sen and the 914 pathways, as well as those of Fer-res and the pathways. We found most of the 14 pathways that were specifically upregulated in DS2 were the most highly correlated with Fer-res, while they were somewhat uncorrelated with Fer-sen (Fig. S9C,D).

### Pathway activity analysis in time course

Of the 351 conditions (compounds and doses) that were tested at all eight time points, 71 were assigned to DS2 and 58 to DSs5-9 on day 29 (Fig. 6A). They were named as ‘tolerance’ and ‘tissue injury’, respectively. The other 222 conditions were neither, and named as ‘others’. At each time point, we assessed whether pathways were changed in tolerance or tissue injury. We computed KS statistics of each pathway per condition, and assessed whether the statistics were deviated between the classes (tolerance vs others; tissue injury vs others) using two-sample two-sided Wilcoxon test (Fig. 6C).

**Fig 6.**
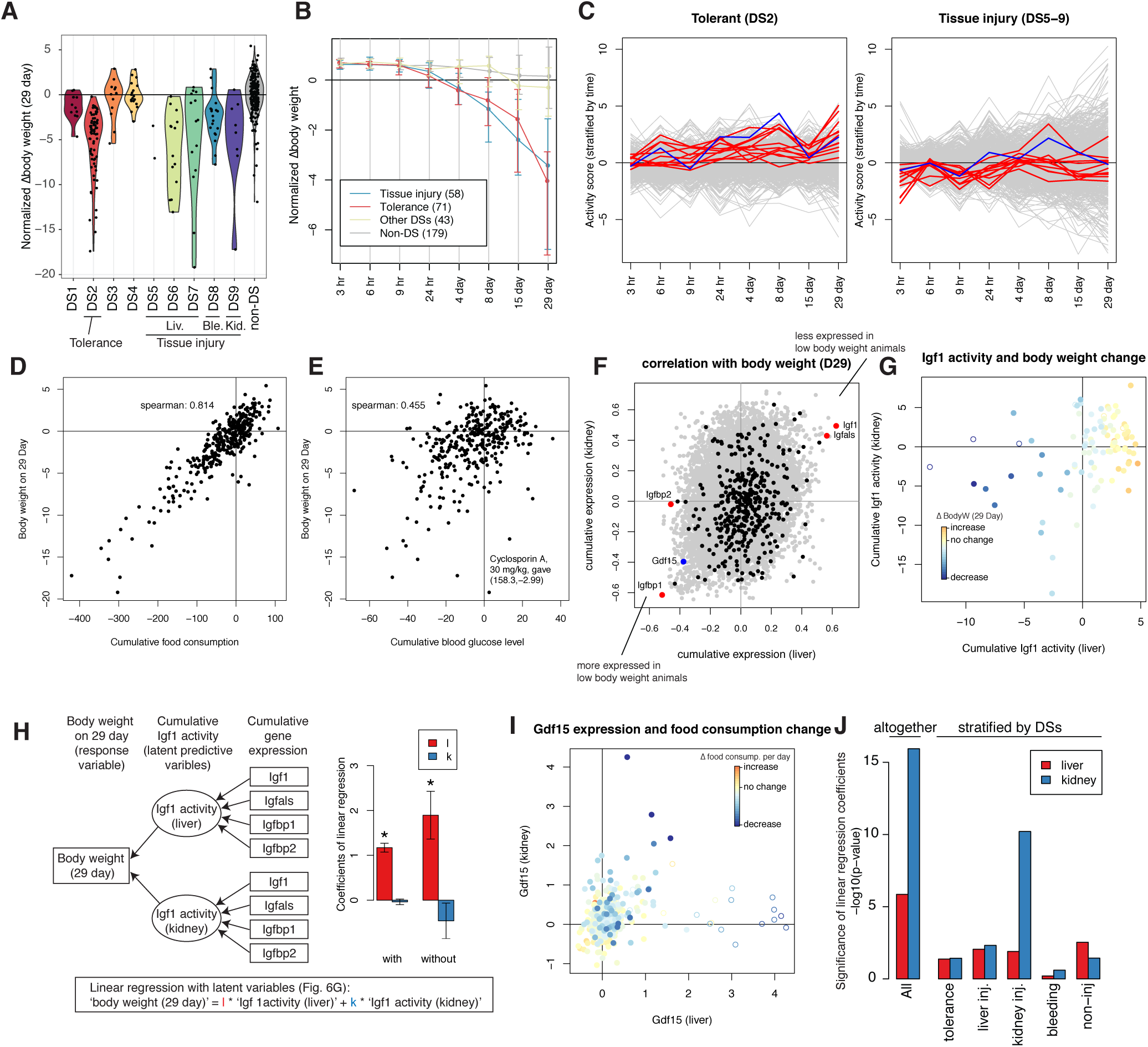
Low blood glucose level inhibits Igf1 paracrine to decrease body weight. **(A)** Changes in normalized body weight on 29 day. **(B)** Body weight change over time. Treatment conditions were grouped into the DSs on 29 day: tolerance (DS2), tissue injury (DSs5-9), other DS (DSs1,3,4), and non-DS. **(C)** Pathway activities in tolerance (71) or tissue injury associated conditions (58) compared with the rest of the conditions (222). Y axis shows relative pathway activities at a given time point (signed log p-values from two-sample Wilcoxon test). Red and blue lines correspond to the 14 DS2-associated pathways listed in Fig. S9D and ‘ferroptosis resistant signature’, respectively. **(D-E)** Relationship between **(D)** cumulative food consumption or **(E)** cumulative blood glucose level and body weight on 29 day; cyclosporine A at 30 mg/kg inducing hyperglycema continuously,therefore became an outlier in **(E). (F)** Spearman correlated with cumulative gene expression in liver and kidney and body weight on 29 day. Plasma proteome genes, four Igf-related genes, and Gdf15 were highlighted in black, red, and blue, respectively. **(G)** Cumulative Igf1 activity over 29 days in liver and kidney, computed in the model Fig. 6H. Points were color coded by the extent of body weight change. Open circles indicate synthetic hormone treatments. **(H)** Linear regression model with latent variables (left). Rectangle and ellipsoidal nodes indicate observed data and latent variables, respectively. Contribution of liver and kidney Igf1 activities to body weight on 29 day, both with and without synthetic hormone data. **(I)** Gdf15 expression in liver and kidney at five time points (1,4,8,15 29 days). Points were color coded by changes in food consumption per day. Open circles indicate synthetic hormone treatments. **(J)** Significance of contributions of liver and kidney Gdf15 expressions to changes in food consumption, computed by linear regression with and without stratification by disease states. See also Figure S10.

### Compiling blood plasma proteins

We assembled the collection of experimentally validated plasma proteins from two different databases. First, from Human Protein Atlas (https://www.proteinatlas.org/), 3704 ‘predicted secretory proteins’ have evidences at protein levels were looked at, 2960 of which have rat orthologs. Second, from Plasma Proteome Database (plasmaproteomedatabase.org/), 468 proteins have more than one associated reference that they were observed in plasma, 382 of which have rat orthologs. Altogether, 376 rat orthologs (unique Entrez gene IDs) were observed from the two databases that were also measured in Affymetrix Rat Genome 230 2.0 Array.

### Correlation between gene expression and food consumption or body weight

We computed area under curve (weighted sum) of the all genes’ expression in liver and kidney over 29 days, including 376 plasma protein-encoding genes for 351 conditions tested at all time points. Then we calculated the Spearman correlation coefficients between the cumulative gene expression and the body weight on 29 day (Fig. 6F). The positive and negative correlation indicates genes were expressed less or more in the animals with decreased body weight. We also computed Spearman correlation coefficients between individual gene expression, not cumulative, and food consumption per day. In which, measurements at five time points (1,4,8,15,29 day) were treated as independent conditions, and correlation was calculated among all conditions and time points (Fig. S10C).

### Linear regression of body weight or food consumption on Gdf15 or Igf1 activities

#### Igf1 and body weight

To assess the relationship between Igf1 and body weight, Igf1 activities in liver and kidney was summarized from four Igf1-related genes in the tissues as latent variables, which were further used to regress changes in body weight on 29 day (as modeled in Fig. 6H). This latent variable analysis was performed using lavaan package. In the model, “Igf1 activity” was conceived as a latent variable each for liver and kidney, which is estimated from cumulative expression of four Igf1-related genes (Igf1, Igfals, Igfbp1, Igfbp2). And the two latent variables, Igf1 activity for liver and kidney, were used to see their contributions in the change in body weight on 29 day.

#### Gdf15 and food consumption

Multivariate linear regression of food consumption on Gdf15 expression in liver and kidney was performed by lm() function. Since Gdf15 level in the tissues are substantially different among DSs (Fig. S10J), the linear regression was also performed with the data stratified into DSs, where DSs were incorporated as a categorical interaction term in lm() (Fig. S10K,L). We converted the significance of the coefficients by computing and plotting signed log p-values (Fig. 6J).

## Results

### Unsupervised identification of disease states from physiology and histology data

There is currently no standard method to classify toxin-induced pathology in a completely automated fashion. Therefore, we explored the physiology and histology space of an existing large toxicogenomic dataset (Open TG-GATEs), using advanced machine learning techniques to identify disease states (Fig. 1A; an overview of the data and our analysis workflow are summarized in Figure S1). We performed an initial characterization of disease states using blood chemistry and body/tissue weight data (hereafter referred to as physiology data), which are unbiased measures corresponding to standard clinical tests used for diagnosis in patients. The dataset includes physiology data for 3,564 total conditions, representing administration of 160 chemicals at three dose levels each, with data collected at eight time points ranging from 3 hr to 29 day (Fig. S2). To highlight dis-/similarity and partial dependencies between 37 physiology parameters across all 3,564 treatment conditions, we visualized these conditions using t-distributed stochastic neighbor embedding (t-SNE), which has recently gained popularity in biology, due to its success in dissecting heterogeneity of conditions from high dimensional correlated data *[17]*. The t-SNE physiology map revealed one large island, which may correspond to normal physiology, and several small islands of abnormal physiology (Fig. 1B, Fig. S3). Naming of these islands, and their relationship to standard toxin-induced pathology states, is discussed below.

We next focused on liver and kidney histology data to ask if they support the discrete disease states that emerged from physiology data. Histology is typically employed after physiology to diagnose human disease. It is also part of the routine exercise required in regulatory toxicity assessments. In Open TG-GATEs, these data were recorded as calls from a standard constrained terminology made by expert toxicologic pathologists based on microscopic examinations of H&E stained tissue sections. One compound treatment can induce more than one histopathology phenotype concomitantly, and typically, a tissue experiencing severe injury exhibits multiple histological phenotypes (Fig. S4–5). We computed ‘severity scores’ based on the number of abnormal histology phenotypes scored. We then color coded the physiology t-SNE map by liver and kidney histopathology severity scores (Fig. 1C). Visual inspection confirmed good correlation between physiology and histology, where small distinct islands exhibited abnormal liver and kidney histology. However, conditions exhibiting similar physiological abnormality did not always agree on the histopathological severity scores. We suspected that this is in part due to lack of reliable severity metrics in histopathology; histopathology calls made by human experts are intrinsically less quantitative than physiology metrics (e.g., low-level changes may be missed) or may suffer from bias and/or inaccuracy due to scoring only a few tissue slices per animal. To resolve such discrepancy between physiology and histopathology, and we decided to primarily rely on physiology rather than histopathology data where there is a disagreement; by imputing the liver and kidney severity scores on the physiology map, we have a more consistency between the two data and we can set unified criteria for identifying pathological conditions relying on both physiology and pathology datasets were set. (Fig. 1D).

We next called disease states by identifying discrete clusters in the conjoined map of physiology and histopathology metrics using a density-based clustering algorithm *[18]*. We iterated the whole procedure from t-SNE to cluster calling 100 times to compensate for the stochastic nature of the t-SNE algorithm *[19]*. Nine consensus clusters emerged robustly across the 100 runs and were termed “disease state (DS)” clusters (Fig. 1E). While use of the word “disease” implies a negative health impact, it is in principle possible that some “disease states” are beneficial. Of 3,564 conditions, each DS contained 37 to 203 conditions. 2,723 conditions did not belong to any DSs, and we classified these as non-disease state (non-DS) (Table S1). DSs are listed in Table 1.

**Table 1.** Characteristics of the nine disease states

| DS | Clinical description | Physiology | Histopathology | Induced transcriptome | Toxin class | Kinetics |
| --- | --- | --- | --- | --- | --- | --- |
| 1 | Metabolically active | Rel.Liver.Weight up | Granular eosinophilic degeneration (L) | Anti-inflammatory; anabolism of ATP, lipids, amino acids (L) | Lipid-lowering drugs | Late-onset; robust |
| 2 | Tolerance | Body.Weight, GLC down | Hypertrophy (L) | Xenobiotic metabolism, ferroptosis resistance (L) | Synthetic hormones | Late-onset; robust |
| 3 | Hemolytic anemia | TBIL up | N/A | N/A | N/A | Early-onset; transient |
| 4 | N/A | TBIL, DBIL down | N/A | N/A | N/A | Early-onset; transient |
| 5 | Acute liver injury | PT up; TC, PL down | Eosinophilic change (L) | IL-1/TNFα signal, RNA-pol II transcription (L) | N/A | Early-onset; transient |
| 6 | Cholesatsis | DBIL up | Prolifeartion of bile ducts (L) | IL-1/TNFα signal, cancer signature, response to alkaloid (L) | N/A | Early-onset; transient |
| 7 | Hepatocellular injury | AST, ALT, LDH up | Single cell necrosis (L) | IL-1/TNFα signal, cancer signature, cell cycle, collagen biosynthesis (L) | N/A | Early-onset; transient |
| 8 | Bleeding | WBC, Ret, Neu up; Hb, Ht, Lym, RBC down | Extramedullary hematopoiesis (L) | IL-1/TNFα signal, IL-6/Jak/STAT signal, complement/coagulation cascade (L) | NSAIDs | Late-onset; robust |
| 9 | Kidney injury | BUN, Rel.Kidney.Weight, up | Hypertrophy (K) | IL-1/TNFα signal (K) | N/A | Late-onset; robust |

This table summarizes the analysis of physiology (Fig. 2B), histology (Fig. 2C), transcriptome (Fig. 3), toxin class (Fig. S6), and kinetics (Fig. 4). Physiology acronyms are described in Fig. S2A. Refer to the corresponding figures for the details.

### Known and novel disease states identified by unsupervised analysis

We next tried to map DSs onto standard pathologies in the toxicology literature, referring primarily to the physiology and histology markers, and in some cases also to subsequent analysis of gene expression changes or toxin mechanisms (described below). Comparison of histopathological severity scores showed which tissues were more affected in each DS (Fig. 2A). A simplified clustergram highlighted physiology parameters that most strongly defined each state (Fig. 2B). As we hoped, a distinct set of histology phenotypes were associated with DSs (Fig. 2C).

Table 1 lists the 9 DSs, naming each one based on the most characteristic change in physiology and/or histopathology, together with transcriptome and kinetics analysis that are described below. DS1-4 were not associated with detectable tissue injury, and would likely elude a conventional toxicogenomic analysis. DS5-9 showed clear signals of tissue injury as scored by physiology and histology, and map clearly onto the kinds of disease used as pre-determined classifiers in conventional toxicogenomics. Recognition of DS1-4 therefore represents a preliminary success of our analysis, and DS2 in particular emerged as mechanistically significant in subsequent analysis. DS1 showed activated synthesis of various metabolites and increased relative liver mass. DS2 represents acquired toxin-induced tolerance while decreasing body weight. In DS3, total bilirubin (TBIL) increased, but direct bilirubin (DBIL), a liver injury marker, did not. Based on this profile we suspect that DS3 corresponds to hemolytic anemia. In DS4, TBIL level was decreased, which is not seen in known clinical states.

DS4 lacked common histological or transcriptome changes in liver and kidney, and may not be a disease state in the conventional sense, though it clearly is abnormal *[20]* and reproducible. DS5-7 were marked by an increase in standard liver injury markers that are used in human diagnosis of liver damage *[21]*. DS5 exhibited longer prothrombin time (PT), suggesting a decrease in the synthesis of prothrombin, a liver-synthesized blood coagulation protein. DS5 would not clinically be considered to be an explicit liver injury, unlike DS6-7, but the data strongly suggest that the liver’s health is affected. The DS6 profile corresponds to cholestasis, an injury of the liver bile ducts. DS7 corresponds to hepatocellular injury. In DS8, multiple hematological parameters were changed and unique histological and transcriptional phenotypes were observed, indicating that the animals suffered from bleeding, induction of synthesis of complement factors and coagulation cascade components, and hematopoiesis in the liver *[22]*. DS9 corresponds to kidney injury. The database contained fewer reference kidney toxins than liver toxins, perhaps explaining why we observed only one disease state that mapped to kidney pathology *[23]* in the analysis.

### Overrepresentation of drug classes in disease states

We next asked whether specific toxin classes reliably induced specific DSs, defining classes as containing multiple compounds with known overlapping biological activity, as summarized in Figure S1C. We found that lipid-lowering drugs mapped to DS1, synthetic hormones mapped to DS2, and non-steroidal anti-inflammatory drugs (NSAIDs) mapped on DS8, each at higher doses, and each more than expected by chance (Table 1, Fig. S6A). NSAIDs are known to cause intestinal bleeding at high doses *[24,25]*, which likely accounts for their mapping to DS8 (Fig. S6B,C). The four lipid-lowering drugs mapped onto DS1 were peroxisomal proliferator-activated receptor alpha (PPAR*α*) agonists (clofibate, fenofibrate, WY-14643) and a cholesterol synthesis inhibitor (simvastatin). PPAR*α* agonists increased peroxisomes, which were recognized as eosinophilic granules in the cells in DS1 *[26]* (Fig. S6D,E). This class of drugs has been well studied in rats, and its lipid-lowering effect in the short term is often described as beneficial rather than pathological. However, we decided to keep our initial notation of “disease states” even for DS1; long term treatment with these drugs frequently causes liver cancers, possibly due to hyperactivation of metabolism to increase the liver biomass *[27]*. The NSAIDs- and fibrates-induced disease states of liver were previously characterized by transcriptome-centric analysis of Open TG-GATEs data *[28]*. Compared to the two drug classes that strongly induced representative physiology and histology phenotypes of the DSs, the effect of synthetic hormones on liver was modest, and lacked conspicuous physiology or histology changes such as hypertrophy (Fig. S6F,G).

### Transcriptome description of disease states

To test if each DS was transcriptionally distinct, we performed elastic net classification of liver and kidney transcriptome data (Fig. S1, Fig. S7A). This generated classifiers that attempted to distinguish conditions assigned to each DS from all the rest of conditions using the liver or kidney transcriptome. All classifiers were found to be powerful for separating DSs (all areas under ROC curves were above 0.85, compared to 0.46–0.63 for randomly drawn samples of the same sizes). Thus, each DS has a characteristic transcriptome both in liver and kidney. To examine the functional implications of DS-specific transcriptional states, we assessed whether 914 GO and KEGG pathways changed their activities in DSs, compared to pooled non-DS clusters. We computed “activity scores” by modifying the standard gene set enrichment analysis (GSEA) to capture the significance of systematic changes of pathway activity among conditions assigned to each DS (See Methods for the detail). We interpreted a large positive or negative activity to indicate that a pathway is substantially up or downregulated compared to corresponding vehicle treatments in the DS. Through this analysis, we found that six DSs (DS1,2,5-8) induced systematic changes of pathway activities in the liver transcriptome; in DS9, changes of pathway activities were observed in the common kidney transcriptome but not in the liver transcriptome; and DS3-4 did not capture any common pathway changes either in liver or kidney (Fig. 3A). Hierarchical clustering of pathway activity in both liver and kidney primarily divided nine DSs into two groups (tissue injury or not) (Fig. 3B); consistently, five DSs associated with tissue injury activated pro-inflammatory cytokine signaling, namely cellular response to proinflammatory cytokines (TNF, IFN-*γ*, IL-1) in the corresponding tissues (Fig. S7B).

Liver was more transcriptionally responsive to xenobiotic stimuli than kidney across many conditions that induced a DS. This is expected because the liver is primarily responsible for detoxifying xenobiotics, although the choice of toxins in the dataset may also over-emphasize the role of the liver. The kidney only exhibited major transcriptional changes in DS9, which also phenotypically corresponded to kidney damage in the physiology and histopathology (Table 1).

To better understand the changes induced by toxin exposure in the six DSs we identified as significantly affecting the liver transcriptome (DS1,2,5-8), we classified the modulated pathways based on their transcriptional activity. Pathways that were transcriptionally activated or suppressed in each DS were mapped onto corresponding nodes on a dendrogram (Fig. 3C, Fig. S7C-E). This allowed us to determine in which contexts the pathways were up-or down-regulated. For example, in all of these six DSs, the liver activated transcription of “cytosolic large ribosomal subunit”; in disease states that reported on tissue injuries (DS5-8) pathways categorized as “cellular response to IL-1 and TNF” was activated in the liver. On the other hand, some pathways such as “p53 signaling” or “collagen biosynthetic process” were activated only in specific states (e.g. hepatocellular injury (DS7)), but not in the other DSs.

### DS2 is a state of drug-induced tolerance

A strength of the Open TG-GATEs data is the collection of multiple time points at constant toxin exposure. Kinetic analysis showed that some DSs are mostly or exclusively late-onset, i.e. after 24hrs (DS1,2,8). Others are early-onset or had no particular trends in kinetics (Fig. 4A, Fig. S8A). Of 365 conditions (compound and dose) whose data were collected at all eight time points, 246 (67%) caused at least one DS at some point, and 90 (25%) caused more than one DS over time (Fig. S8B).

To inform on causal connections between DSs we focused on conditions causing more than one DS, and classified temporal transitions at constant toxin exposure (Fig. 4B). This analysis revealed bi-directional inter-conversion between different DSs corresponding to liver damage, which may be expected given transcriptional and histologic overlaps between these related DSs. A strong, unexpected feature was conversion of multiple liver and kidney DSs to DS2. Since DS2 is not associated with liver or kidney injury by histology, this suggests a time-dependent reduction of organ-specific pathology. Fig 4C shows temporal changes in a standard liver injury biomarker (AST), a standard kidney injury biomarker (BUN) and a representative biomarker of DS4 (TBIL) across the transitions out of other DSs and into DS2. Transitions into DS2 were accompanied by reduced injury biomarkers in most cases. After examining the transcriptome data on the different DSs (see below), we interpret this as evidence of induced tolerance.

Toxin–induced tolerance to toxin action, also known as autoprotection *[29,30]*, is poorly understood but is presumably an important component of how xenobiotic resistance evolved, and how modern vertebrates adapt to toxic environments. Analysis of dynamics of DSs and injury biomarkers (Fig. 4B,C) suggests that DS2 is a state of induced tolerance. To determine molecular mechanisms that might drive tolerance and result in DS2, we first re-examined genes that are selectively regulated in this DS. Xenobiotic catabolism genes were strongly and selectively induced in DS2 compared to all other DS and non-DS conditions (Fig. 3D). While this finding might be expected, it emphasizes the function of xenobiotic defense genes, and supports our characterization of DS2 as a state of tolerance.

### Tolerance is associated with induced resistance to ferroptosis

Xenobiotic metabolism is a multi-step reaction: in phase I, cytochrome P450 monooxygenases (Cyp450) conjugate xenobiotic compounds with oxygen using NADPH; in phase II, the products of this reaction are conjugated with hydrophilic groups such as sugars (e.g. glucuronic acid) and glutathione (GSH) to facilitate excretion. When we regrouped xenobiotic catabolism genes based on co-factors, we found that genes encoding NADPH- and GSH-utilizing enzymes were among the most highly expressed genes in DS2 (Fig. 5A, Fig. S9A). Activation of redox metabolism functions is also involved in ferroptosis, a form of cell death that has been implicated in drug-induced liver injury *[31]*. Thus, we also suspected that the liver acquires resistance to ferroptosis in DS2.

Ferroptosis occurs when activity of the selenoprotein glutathione peroxidase 4 (Gpx4) is inhibited (Fig 5B) *[32]*. Ferroptosis pathways are not represented in current GO terms, in part because genes that regulate ferroptosis are still poorly understood *[16]*. To generate an objective classifier of ferroptosis sensitivity, we referred to our previous pharmacogenomic analysis of the NCI-60 project, where we contrasted cellular responses to chemicals known to induce cell death via a number of mechanisms, including ferroptosis (Fer), DNA damage (DNA) and tyrosine kinase inhibition (TKI) in the NCI-60 human cancer cell line panel *[33]*. Using those data, we generated gene expression signatures that serve as markers for sensitivity (sen) or resistance (res) for each of these treatments. We converted these six signatures to rat orthologs and evaluated them across all nine DS compared to non-disease samples. Fer-res was strongly and uniquely upregulated by DS2 (Fig. 5C). Conversely, DSs associated with liver injuries downregulated Fer-res and upregulated Fer-sen, consistent with the role of ferroptosis in liver injury. DS2 does not increase DNA-res or TKI-res, suggesting that toxin-induced tolerance is associated with acquiring resistance to ferroptosis, but not to other cell death mechanisms.

There was no substantial overlap between genes involved in Fer-res and the 14 pathways activated exclusively in DS2 (Fig. S9B). Nevertheless, the transcriptional activity of Fer-res among 3,528 liver transcriptome is most highly correlated with the scores of the 10 GO and KEGG pathways exclusively upregulated in DS2 while the activity of Fer-sen is somewhat anticorrelated with them (Fig. S9C,D). Our analysis supports the hypothesis that DS2 not only overexpresses xenobiotic metabolism genes but also acquires resistance to ferroptosis.

### Whole body response to toxins

Toxin responses have a whole body impacts in addition to organ-specific effects, but these have been much less studied in the toxicology literature, or the human liver/kidney disease literature. Loss of body weight is a well known indicator of chronic toxicity in rats, though its etiology is poorly understood. It likely corresponds to cachexia phenotypes in man, which are well-recognized contributors to disease burden *[34]*. Body weight decrease was the most significant physiological descriptor for DS2, but it was not unique to DS2: all DSs associated with tissue injury (DS5-9) exhibited body weight decrease to a similar extent to DS2 (Fig. 6A,B), while tolerance-associated pathways were activated in DS2, but not in DS5-9, highlighting the differences between the states (Fig. 6C). Toxin-induced weight loss could result from direct responses of peripheral tissues such as muscle and adipose tissues to toxin, or (more likely) from their indirect responses to signals from toxin-exposed tissues that alter global metabolism or/and feeding behavior. The liver and kidney communicate with the whole body through concentrations of multiple metabolites in blood, and also through secreted signaling proteins. We therefore analyzed metabolite and secretome changes in conditions that caused body weight loss, irrespective of whether tolerance was induced.

Suppression of food intake over time was strongly correlated with body weight, as might be expected (Fig. 6D). Food consumption is decreased in conditions associated with tolerance (DS2) or tissue injury (DS5-9) (Fig. S10A), and this may therefore be part of the mechanism leading to loss of body weight in these conditions. We observed a weaker correlation between blood glucose level and weight loss (Fig. S10B, Fig. 6E). The causal chain here appears to be that decreased food consumption weakly correlates with decreased glucose concentration, and decreasing blood glucose is known to cause decreased body weight.

To identify candidate secreted proteins that mediate whole organism toxin responses, we computed cumulative gene expression over 29 days for all genes that changed expression in liver and kidney respectively, and calculated the Spearman correlation between their expression and the body weight on Day 29 (Fig. 6F). To identify secreted proteins we referred to a list of 376 blood plasma proteins measured by proteomics and other methods *[35,36]*. We discovered that the four most strongly up- and down-regulated secreted proteins in the liver were all related to insulin-like growth factor-1 (Igf-1): low body weight animals consistently upregulated the Igf-1 antagonists Igfbp1 and Igfbp2, and downregulated Igf1 itself and its activator Igfals (Fig. 6F) *[37]*. Three of the four proteins showed similar trends in the kidney. We also found that the Spearman correlations of gene expression levels and food consumption across five time points (1, 4, 8, 15, 29 days) showed quite similar results (Fig. S10C), strongly indicating that Igf1 signaling decreased upon suppression of food consumption. These four proteins thus appear to collectively mediate strong organ-to-body communication as part of toxin responses. Igf1 promotes tissue growth *[38]*, so decreasing its activity via gene expression may cause body weight decrease in toxin-exposed rats. Igf-1 secretion also responds to blood glucose levels, so it might be acting as part of a feedback response *[37,39]*. Both liver and kidney decreased Igf1 signaling, but the liver was found to contribute to body weight loss more significantly (Fig. 6G-H, Fig. S10D-F). This is consistent with the fact that Igf1 synthesized and secreted by the liver accounts for 75-80% of paracrine Igf1 found in the blood plasma*[37]*.

Another endocrine factor that might play a significant role in body weight decrease is Gdf15, whose cumulative expressions both in liver and kidney were significantly negatively correlated with body weight (Spearman < −0.2, p-value < 4.9e-7) (Fig. 6F). This TGF*β* family member can be produced by many tissues, and is known to be a toxin response gene [40–42]. It negatively regulates feeding by binding to the receptor Gfral in the brainstem [43–46]. Animals upregulating Gdf15 mRNA either in liver or kidney decreased body weight (Fig 6I, Fig. S10H,I), consistent with a role of Gdf15 in suppressing appetite. However, high expression of Gdf15 mRNA in the liver was induced only by synthetic hormones such as ethinylestardiol or tamoxifen, which induced decrease in body weight without any physiological or histopathological signs of toxicity (Fig. S6F,G). We therefore concluded that they were causing weight loss via a mechanism that differs from classic toxin responses. When these compounds were removed, we found that Gdf15 expression in the kidney, and not the liver, responded consistently to tissue injury (Fig. 6J, Fig. S10J-L). Thus, Gdf15 expression by the kidney after tissue injury may be the general mechanism for toxin-induced weight loss, via negative regulation of feeding. Fig 7 summarizes our current view of the role of liver-body communication in toxin-induced body weight decrease, where liver-to-whole body signals that decrease feeding remain to be identified.

**Fig 7.**
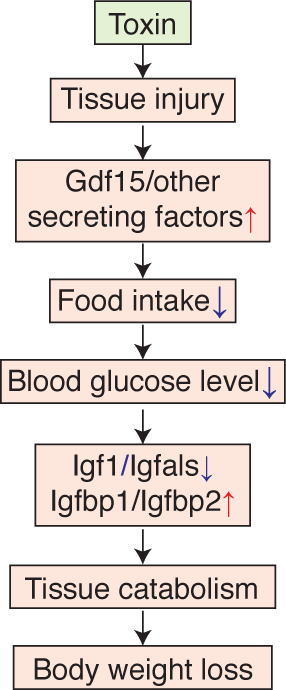
Proposed mechanism of drug-induced body weight loss. (1) Toxin induce tissue injury. (2) Injured tissue and the kidney secrete anorexic factors including Gdf15. (3) Hypothalamus suppresses appetite and food consumption. (4) Blood glucose concentration is decreased. (5) Production of paracrine Igf1 is decreased in the liver. (6) Tissues, such as muscle and adipose, catabolize themselves due to suppression of glucose uptake. (7) Body weight is decreased.

## Discussion

Application of machine learning to physiology and histology data is an emerging field with high potential in translational research *[47]*. We applied a fairly simple unsupervised characterization approach to discover disease states in a high-quality toxicogenomics dataset. Physiology data alone identified nine discrete disease states (Fig. 1). Histopathology data supported these states with high statistical confidence, and helped us determine their relevance to conventional toxin-induced pathologies (Fig. 2D). However, our method required imputing missing phenotypes to compensate for lack of quantitative metrics and likely missing values, potentially including sampling error, in the histopathology calls made by human experts. Open TG-GATEs includes large H&E images of most conditions, and an interesting future question is whether application of artificial intelligence machine vision approaches could help boost reliability and quantification of histopathology images.

Our approach provides a new window into the biology present in a toxicogenomic dataset, but we did not use it to try to improve predictive toxicology. That important goal might be feasible in future studies. Our immediate interest was in how well an algorithm can mimic a physician in terms of defining disease states, which is generally relevant to applications of machine learning in medicine. We were also interested in how the liver, and the organism as a whole, respond, and in some cases adapt, to continuous toxin exposure. Toxin-induced tolerance is highly relevant to environmental and pharmacologic toxicology, and to the evolution of xenobiotic defenses, but has received relatively little attention in the genomic era. Our DS analysis (Table 1) was successful in identification of standard organ pathology states used in toxicology. It also identified several non-standard states with interesting physiology, which demonstrates the potential of computational analysis in translational research and medicine.

An unexpected and interesting outcome of DS kinetics was evidence for acquisition of tolerance to xenobiotics by the liver, also known as autoprotection or pharmacokinetic tolerance. We identified DS2 as the main tolerance state. We then showed, using gene expression analysis, that tolerance is achieved by expected mechanisms that include overexpressing xenobiotic catabolizing enzymes, particularly NADPH- and GSH-dependent ones. Unexpectedly, we found that tolerance correlated with induction of biomarkers for ferroptosis resistance, using an independent dataset of cancer cells responds to drugs to identify these biomarkers. Overall, our analysis of tolerance confirms the expected role of conventional detoxification enzymes, and points to ferroptosis resistance as a novel mechanism worthy of additional study.

Finally, we used transcriptomics to analyze possible molecular causes of body weight decrease, and the role of secretory plasma proteins in orchestrating it. Despite the universal use of body weight as a biomarker of toxicity, its molecular basis is poorly understood. Body weight decrease was found in all DSs where organ injury was present, and also in the tolerance state DS2. As expected, it correlated with decreased food intake and blood glucose (Fig. 6D,E). Remarkably, the four liver-secreted proteins that correlated best with body weight decrease were all part of the Igf1 system, and all changed in the same direction, of decreased Igf1 signaling in disease states associated with weight loss (Fig. 6F). Their high correlation suggests Igf1 shutdown is a common response to diverse tissue injuries caused by toxins, where the chain of causality likely proceeded from decreased feeding causing low blood glucose levels which then causes decreased Igf1 signaling *[37]*. Paracrine Igf1 signaling serves to increase muscle and adipose growth, so its loss may plausibly drive muscle atrophy and body weight loss (Fig. 7). This leaves open the question of how organ damage triggers decreased feeding. Gdf15 is an interesting candidate. It is known as a common early marker of drug-induced liver injury *[41,48]*, and binds to Gfral in the brainstem to suppress food intake [43–46]. Our data strongly support a role of Gdf15 in mediating weight loss in response to multiple toxins (Fig. 6I,J, Fig. S10J,L). The data also suggest that Gdf15 in response to multiple toxins are primarily synthesized in kidney, and most strongly induced by kidney toxins, consistent with a recent report that Gfral mediates weight loss in response to the kidney toxin Cisplatin *[46]*. However, Gdf15 expression by liver was only strongly correlated with weight loss in response to certain synthetic hormones that were not overt toxins (Fig 6I,J, Fig. S10H,J). Therefore, as yet unidentified signals are more likely to mediate liver-to-whole-body signaling to suppress food intake upon synthetic hormone treatments.

In summary, our approach shows the value of systematic data collection and analysis for revealing new organ biology. Ferroptosis emerged as an important factor in toxin responses, and may be a druggable driver of tissue pathophysiology. We made progress on the little-studied problem of how organ toxicity triggers whole body weight decrease, which is relevant to mechanistic toxicology, effects of alcohol in man, and perhaps to diagnosis and treatment of human diseases with non-toxin causes. From a translational perspective, our findings support a role of GDF15 antagonists for treatment of drug-induced cachexia. They particularly support a role of IGF1 agonists for treating cachexia more universally. Their use is complicated by glucose-lowering side effects, but we suggest they deserve more attention for treatment of cachexia across multiple diseases. Igf1 modifiers that are co-regulated during toxin-induced weight loss (Igfbp1, Igfbp2 and Igfals) are also worthy of consideration as druggable targets in cachexia.

## Acknowledgments

We thank Rebecca Ward, Laura Maliszewski, Debora Marks, and Peter Koch of Harvard Medical School, Scott Dixon of Stanford University, Etsuko Ohta and Kenji Kubara of Eisai Tsukuba Research Laboratories, and Dahlene Fusco of Massachusetts General Hospital for helpful discussion.

## Author contributions

K.S.: Conceptualization, Formal Analysis, Investigation, Data Curation, Writing, Visualization, Project Administration. T.J.M.: Conceptualization, Writing, Supervision.

## Data and materials availability

Original data are publicly available on Open TG-GATEs website (http://toxico.nibiohn.go.jp/open-tggates) and Life Science Database Archive (https://dbarchive.biosciencedbc.jp/en/open-tggates). Processed data in this study are available as supplemental Information of this paper, from the following link: https://figshare.com/s/7fe89f0f11aada6ba7df

## Supplemental Information

Figure S1. Experimental and analytical workflow of this study

Figure S2. Physiological parameters and their correlations

Figure S3. Physiological parameters on t-SNE map

Figure S4. Kinetics of histopathology characterization

Figure S5. Histopathology mapped onto the physiology t-SNE map

Figure S6. Toxin class overrepresentations in disease states

Figure S7. Transcriptome analysis for DS classification and kinetics

Figure S8. Disease state dynamics

Figure S9. Induced drug tolerance partly due to resistance to ferroptosis

Figure S10. Mechanism of body weight loss

Supplemental Data 1. Treatment conditions

Supplemental Data 2. Physiology parameters of 3,564 conditions

Supplemental Data 3. Histopathology parameters of 3,564 conditions

Supplemental Data 4. Liver normalized transcriptome of 3,528 conditions

Supplemental Data 5. Kidney normalized transcriptome of 975 conditions

Supplemental Data 6. Conditions assigned to nine DSs and non-DS

Supplemental Data 7. Significantly changed pathways

Supplemental Data 8. Xenobiotic metabolism enzymes transcriptionally activated in DS2

Supplemental Data 9. Food consumption data of 337 conditions over 29 days Supplemental Figures are submitted as a separate pdf file. Supplemental Data are available from the following link: https://figshare.com/s/7fe89f0f11aada6ba7df

